# Activity of a long-acting injectable bedaquiline formulation in a paucibacillary mouse model of latent tuberculosis infection

**DOI:** 10.1101/515692

**Authors:** Amit Kaushik, Nicole C. Ammerman, Sandeep Tyagi, Vikram Saini, Iwan Vervoort, Sophie Lachau-Durand, Eric Nuermberger, Koen Andries

**Affiliations:** Center for Tuberculosis Research, Department of Medicine, Johns Hopkins University School of Medicine, Baltimore, Maryland, USA; Janssen R&D, a division of Janssen Pharmaceutica NV, Beerse, Belgium

## Abstract

The potent anti-tuberculosis activity and long half-life of bedaquiline make it an attractive candidate for long-acting/extended release formulations for treatment of latent tuberculosis infection (LTBI). Our objective was to evaluate a long-acting injectable (LAI) bedaquiline formulation in a validated paucibacillary mouse model of LTBI. Following immunization with *Mycobacterium bovis* rBCG30, BALB/c mice were challenged by aerosol infection with *M. tuberculosis* H37Rv. Treatment began 13 weeks after challenge infection with one of the following regimens: untreated negative control; positive controls of daily rifampin (10 mg/kg), once-weekly rifapentine (15 mg/kg) and isoniazid (50 mg/kg), or daily bedaquiline (25 mg/kg); test regimens of one, two, or three monthly doses of LAI bedaquiline at 160 mg/dose (B_LAI-160_); and test regimens of daily bedaquiline at 2.67 (B_2.67_), 5.33 (B_5.33_), or 8 (B_8_) mg/kg to deliver the same total bedaquiline as one, two, or three doses of B_LAI-160_, respectively. All drugs were administered orally, except for B_LAI-160_ (intramuscular injection). The primary outcome was the decline in *M. tuberculosis* lung CFU counts during 12 weeks of treatment. The negative and positive control regimens performed as expected. One, two, and three doses of B_LA-160_ resulted in decreases of 2.9, 3.2, and 3.5 log_10_ CFU/lung, respectively by week 12. Daily oral dosing with B_2.67_, B_5.33_, and B_8_ decreased lung CFU counts by 1.6, 2.8, and 4.1 log_10_, respectively. One dose of B_LAI-160_ exhibited activity for at least 12 weeks. The sustained activity of B_LAI-160_ indicates promise as a short-course LTBI treatment requiring few patient encounters to ensure treatment completion.

## INTRODUCTION

The crux of global efforts to eliminate tuberculosis (TB) is the prevention of *Mycobacterium tuberculosis* transmission in the population. With approximately 10 million incident cases of TB occurring annually (1), the identification and treatment of individuals with active disease is clearly necessary for interrupting transmission. Addressing incident TB in the population, however, will not be sufficient to end the global epidemic, as the World Health Organization (WHO) has stated that up to one-third of the world’s population may be latently infected with *M. tuberculosis*, referred to as latent TB infection (LTBI) (2). This reservoir of potentially billions of people serves as an ever-present source of new TB cases, independent of recent transmission events (3). Thus, the identification and treatment of individuals with LTBI is also an essential component of the WHO’s End TB Strategy (2, 4).

There are currently four WHO-recommended LTBI treatment regimens: daily isoniazid monotherapy for 6-9 months, daily rifampin for 3-4 months, daily rifampin plus isoniazid for three months, and weekly rifapentine plus isoniazid for three months (2). When completed, all four regimens are highly and equivalently efficacious in reducing the risk of developing active TB disease, but the shorter (*i.e.*, 3-4 month) regimens are associated with higher rates of completion than the longer (*i.e.*, 6-9 month) regimen (5–7). However, ensuring treatment completion of the three- or four-month regimens is still a formidable challenge for TB control programs. The availability of efficacious regimens even shorter than three months could further improve treatment completion rates. An example is the 1-month daily rifapentine plus isoniazid regimen recently evaluated in a phase 3 clinical trial for the treatment of LTBI in individuals infected with HIV (8); the efficacy of this regimen was not inferior to the 9-month control regimen of daily isoniazid and was associated with statistically significantly higher rates of treatment completion.

Beyond decreasing regimen durations, another modification with potential to significantly reduce the burden on healthcare delivery systems and improve LTBI treatment completion rates is the use of long-acting injectable (LAI) formulations for drug administration (9). The development and administration of LAI and implantable drug formulations have improved adherence, *i.e.*, fewer missed “daily” doses following injection, compared to daily oral drug intake in patients receiving anti-psychotic medications (10) and in individuals using hormone-based contraceptives (11). Recently, significant advances have been made in the development of LAI formulations of antiretroviral drugs administered monthly or bi-monthly for both prevention and treatment of HIV infection (12–15), with LAI formulations of cabotegravir and rilpivirine currently being evaluated in a phase 3 randomized clinical trial in adults with HIV-1 infection (ClinicalTrials.gov identifier NCT02951052). LAI formulations may be easier for children than swallowing daily medications [(16) and ClinicalTrials.gov identifier NCT03497676], and importantly, several studies have indicated a high level of interest in and preference for long-acting injectable forms, versus daily oral administration, of HIV pre-exposure prophylaxis across diverse populations (17–20).

Although LAI formulations may be well-suited for use in LTBI treatment, not all anti-TB drugs are well-suited for LAI formulations. Two key properties of (pro)drugs administered in LAI formulations are low aqueous solubility, to preclude rapid dissolution and release of the active drug substance, and a reasonably long pharmacokinetic (PK) elimination half-life, *i.e.*, slow clearance from the body (21). For an antimicrobial, another desired property is high potency, negating the need for high concentrations in the blood (9) and allowing low drug doses to be injected. The diarylquinoline bedaquiline, with a low minimum inhibitory concentration (MIC) for *M. tuberculosis* (about 0.03 μg/mL), high lipophilicity (logP 7.3), and a long half-life (about 24 hours functionally or effectively), possesses a profile that may be suitable for use in an LAI formulation (9, 22–25). Furthermore, bedaquiline has been shown to specifically contribute significant treatment-shortening activity in mouse models of TB (26–28) and is associated with treatment-shortening in patients with multidrug-resistant-(MDR-) TB (29–31), suggesting that this drug could also contribute treatment-shortening activity to an LTBI treatment regimen. We previously demonstrated in a validated paucibacillary mouse model of LTBI that three months of daily, orally-administered bedaquiline had equivalent or superior sterilizing activity compared to each of the four WHO-recommended regimens (32). Thus, an LAI formulation of bedaquiline has the potential to significantly shorten and simplify LTBI treatment, including MDR-LTBI treatment.

Here, we describe the PK and activity of an LAI bedaquiline formulation in the validated paucibacillary mouse model of LTBI (32–34) by comparing the bactericidal activity of one, two, or three monthly injections of the long-acting formulation to that of the same total doses of bedaquiline administered daily by the oral route.

## RESULTS

### MIC of long-acting bedaquiline formulation for *M. tuberculosis* H37Rv

Using a broth macrodilution assay, the MIC of the long-acting bedaquiline microsuspension formulation for *M. tuberculosis* H37Rv was 0.03 μg/mL. This was identical to the MIC of the bedaquiline powder used for oral drug administration, and in agreement with previously published MIC values of bedaquiline for this strain (28, 35).

### PK characteristics of long-acting bedaquiline formulation

The mouse plasma concentration-time profiles of bedaquiline and its M2 metabolite are displayed in Figure 1, and the plasma PK parameters are presented in Table 1. After intramuscular administration of 160 mg/kg bedaquiline microsuspension, the release of bedaquiline from the injection site was slow as shown by plasma concentrations above the MIC at 2184 h (13 weeks) post-dosing. The C_max_ of bedaquiline was reached earlier (range 1-4 h) compared to that of M2 (range 24-168 h). Considering the interindividual variability, the C_max_ values were not significantly different between bedaquiline and M2, whereas the AUC_0-∞_ value for M2 was 3.6-fold higher than that of bedaquiline.

**Figure 1.**
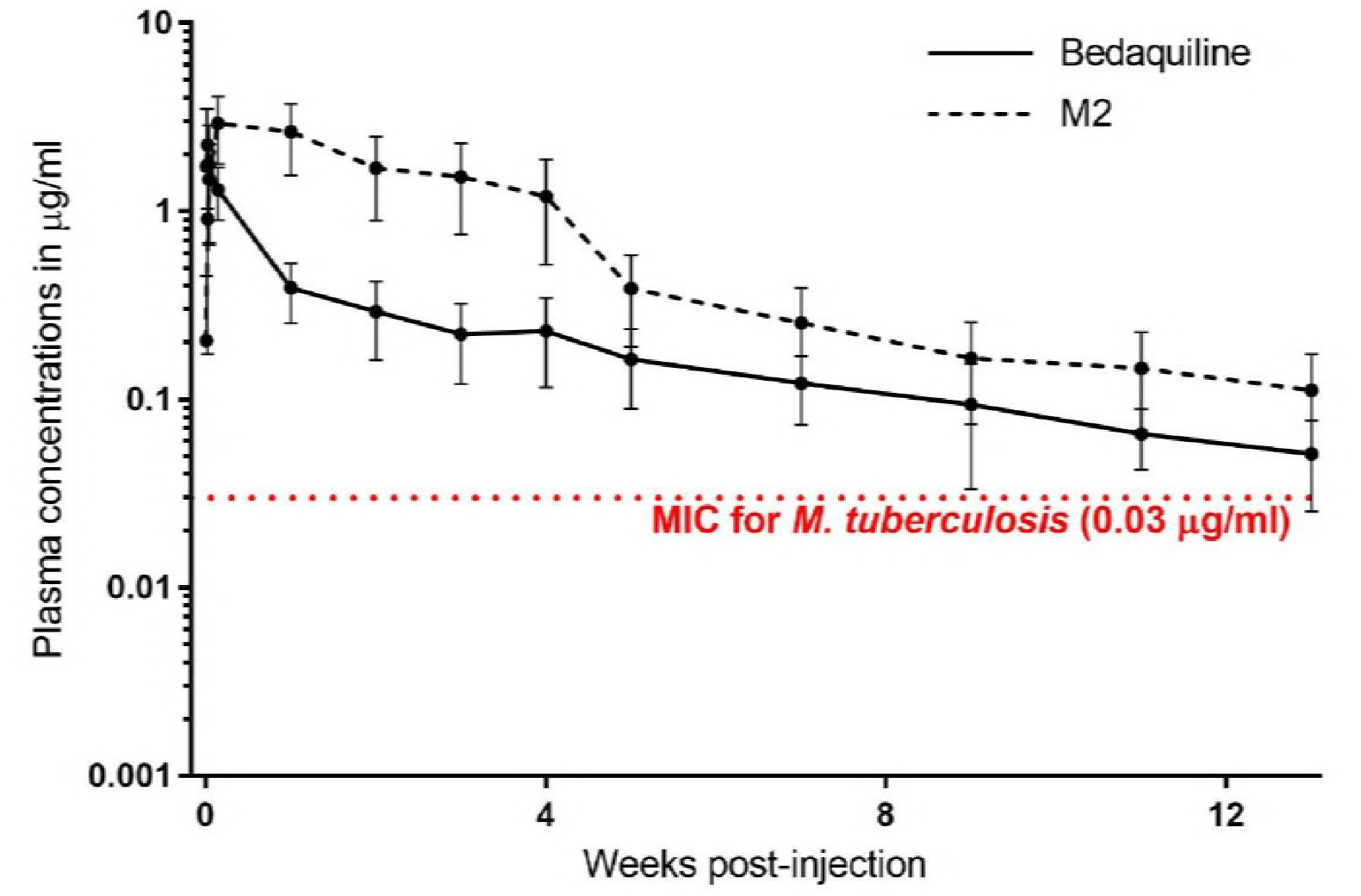
Plasma concentrations of bedaquiline and its M2 metabolite following a single, 160 mg/kg intramuscular injection of long-acting bedaquiline. Data points represent mean values, and error bars represent standard deviation (5 mice sampled per time point).

**Table 1.** Plasma pharmacokinetic parameters of bedaquiline and M2 after a single, 160 mg/kg intramuscular injection of long-acting bedaquiline.

| Pharmacokinetic parameter | Bedaquiline | M2 |
| --- | --- | --- |
| C<sub>max</sub> (ng/mL) | 2,363 (1,447) | 3,002 (1,139) |
| T<sub>max</sub> (h)* | 1-4 | 24-168 |
| AUC<sub>0-2184h</sub> (ng•hr/mL) | 447,361 (174,979) | 1,689,422 (755,169) |
| AUC<sub>0-∞</sub> (ng•hr/mL) | 500,850 (201,928) | 1,823,672 (836,740) |

\*A range is given for T<sub>max</sub> values; all other data represent means (standard deviation) for five mice sampled at each time point.

### Establishment of stable, low-level *M. tuberculosis* infection

Mice were immunized by aerosol infection with *M. bovis* rBCG30; the day after immunization, the mean rBCG30 lung CFU count was 3.05 (SD 0.10) log_10_ (**Tables S1, S2**). Six weeks later, mice were challenged by aerosol infection with *M. tuberculosis* H37Rv. The day after challenge infection, the mean *M. tuberculosis* lung CFU count was 2.11 (SD 0.09) log_10_, and the mean rBCG30 lung CFU count was 4.95 (SD 0.11) log_10_ (**Tables S1, S2**). Thirteen weeks later, on the day of treatment initiation (Day 0), the mean *M. tuberculosis* lung CFU count was 4.75 (SD 0.27), and mean lung CFU counts for the untreated control mice (Table 2) remained at this level throughout the experiment: 4.71 (SD 0.48), 4.60 (SD 0.27) and 4.94 (SD 0.29) log_10_ at Weeks 4, 8, and 12, respectively (Table 3, **Figure S1**, and **Tables S3-S6**). Thus, a stable, relatively low-level *M. tuberculosis* lung infection was established in the mice.

**Table 2.** Regimens evaluated for 12 weeks in a paucibacillary mouse model of LTBI treatment.

| Regimen name | Regimen description |
| --- | --- |
| Untreated | Negative control, no drug administered |
| R<sub>10</sub> (5/7) | Positive control, rifampin (R) at 10 mg/kg, administered daily* by gavage |
| H<sub>50</sub>P<sub>15</sub> (1/7) | Positive control, isoniazid (H) at 50 mg/kg and rifapentine (P) at 15 mg/kg, administered once weekly by gavage** |
| B<sub>25</sub> (5/7) | Positive control, bedaquiline (B) at 25 mg/kg administered daily by gavage |
| B<sub>8</sub> (5/7) | B at 8 mg/kg administered daily by gavage |
| B<sub>5.33</sub> (5/7) | B at 5.33 mg/kg administered daily by gavage |
| B<sub>2.67</sub> (5/7) | B at 2.67 mg/kg administered daily by gavage |
| B<sub>LAI-160</sub> (1/28) × 3 | Long-acting injectable bedaquiline formulation (B<sub>LAI</sub>) at 160 mg/kg, administered every 28 days by intramuscular injection, for three total doses: Day 0, Day 28 (Week 4), and Day 56 (Week 8) |
| B<sub>LAI-160</sub> (1/28) × 2 | B<sub>LAI</sub> at 160 mg/kg, administered every 28 days by intramuscular injection, for two total doses: Day 0 and Day 28 (Week 4) |
| B<sub>LAI-160</sub> (1/28) × 1 | B<sub>LAI</sub> at 160 mg/kg, administered once on Day 0 |

\* “Daily” indicates administration Monday through Friday.
\*\* Isoniazid was administered at least one hour after rifapentine administration. The once weekly gavage occurred every Monday.

**Table 3.** *M. tuberculosis* lung CFU counts.

| Regimen | Total B dose<br>(mg/kg)<br>administered<br>in 12 weeks | Mean (SD) *M. tuberculosis* log<sub>10</sub> CFU/lung at the<br>indicated time point: |  |  |  |  |
| --- | --- | --- | --- | --- | --- | --- |
|  |  | Week -13 | Day 0 | Week 4 | Week 8 | Week 12 |
| Untreated | N/A | 2.11<br>(0.09) | 4.75<br>(0.27) | 4.71<br>(0.48) | 4.60<br>(0.27) | 4.94<br>(0.29) |
| R<sub>10</sub> (5/7) | N/A |  |  | 3.39<br>(0.46) | 2.74<br>(0.62) | 1.27<br>(0.85) |
| H<sub>50</sub>P<sub>15</sub> (1/7) | N/A |  |  | 2.67<br>(0.25) | 0.79<br>(0.80) | 0.28<br>(0.41) |
| B<sub>25</sub> (5/7) | 1500 |  |  | 3.01<br>(0.45) | 0.82<br>(0.49) | 0.07<br>(0.09) |
| B<sub>8</sub> (5/7) | 480 |  |  | 3.30<br>(0.12) | 2.42<br>(0.26) | 0.69<br>(0.43) |
| B<sub>5.33</sub> (5/7) | 320 |  |  | 3.83<br>(0.25) | 3.15<br>(0.47) | 1.98<br>(0.17) |
| B<sub>2.67</sub> (5/7) | 160 |  |  | 3.96<br>(0.35) | 3.52<br>(0.38) | 3.16<br>(0.24) |
| B<sub>LAI-160</sub> (1/28) × 3 | 480 |  |  |  |  | 1.23<br>(0.16) |
| B<sub>LAI-160</sub> (1/28) × 2 | 320 |  |  |  | 2.31<br>(0.40) | 1.63<br>(0.40) |
| B<sub>LAI-160</sub> (1/28) × 1 | 160 |  |  | 3.55<br>(0.32) | 3.31<br>(0.38) | 1.83<br>(0.34) |

### Activity of LAI bedaquiline in a mouse model of LTBI

On Day 0, treatment was initiated with the regimens described in Table 2. Throughout the 12 weeks of treatment, the daily rifampin and the once-weekly rifapentine-isoniazid control regimens performed as expected (32–34), resulting in total reductions of about 3.5 and 4.5 log_10_ CFU/lung, respectively (Table 3, **Figure S1A**, and **Tables S3-S7**). Daily oral dosing with bedaquiline at 25 mg/kg also performed as expected (32, 33), resulting in a reduction of about 4.7 log_10_ CFU/lung over 12 weeks of treatment.

For oral bedaquiline regimens, increasing bactericidal activity was observed with increasing dose at Weeks 4, 8, and 12 (Table 3, **Figure S1B**, and **Tables S3-S7**). After 12 weeks of treatment, only the B_2.67_ (5/7) regimen was significantly less bactericidal than the R_10_ (5/7) control (p < 0.0001). For mice that received one or two injections of B_LAI-160_ (1/28), lung CFU counts were equivalent to those in mice that received the same total bedaquiline dose administered as a daily oral regimen, B_8_ (5/7) for 4 or 8 weeks, respectively (p > 0.05). The lung CFU counts in mice that received a single injection of B_LAI-160_ declined between each time point (**Figure S1C**). After 12 weeks of treatment, their CFU counts were lower than those observed in mice that received the same total dose of bedaquiline (160 mg/kg) via daily oral dosing with B_2.67_ (5/7) (p = 0.0002), with the former regimen resulting in a decline of about 3 log_10_ CFU/lung and the latter resulting in a decline of 1.7 log_10_ CFU/lung, compared to untreated control mice (**Figure S1D**). In mice that received a total bedaquiline dose of 320 mg/kg, either through two injections of B_LAI-160_ or through daily oral dosing of B_5.33_ (5/7), the decline in lung CFU counts was similar at about 3 log_10_ CFU/lung (p > 0.05) (**Figure S1E**). For mice that received a total bedaquiline dose of 480 mg/kg via three injections of B_LAI-160_ (1/28), the lung CFU counts were modestly higher than in mice that received the equivalent total dose through daily oral dosing with B_8_ (5/7) (**Figure S1F**), and the difference was not statistically significant.

## DISCUSSION

Although highly efficacious regimens exist for the treatment of LTBI, low treatment completion rates hinder their practical effectiveness (2, 6, 7, 36). The use of LAI drug formulations for LTBI treatment could simplify treatment and improve completion rates (9). It may also overcome limitations of poor or variable oral bioavailability, mitigate concentration-dependent toxicity and drug-drug interactions, and ease administration in children. A bedaquiline LAI-based LTBI regimen could confer these benefits to contacts of MDR-TB patients, for whom short-course rifamycin-containing regimens are likely to be ineffective as preventive therapy. Here, we report the development and initial evaluation of an LAI formulation of bedaquiline with a promising PK and pharmacodynamic profile for use in LTBI treatment.

While the bacterial burden associated with human LTBI is unknown, it is considered to be less than 4 log_10_ CFU, as the lower limit of detection with acid-fast smears is approximately 4 log_10_ CFU/mL sputum (37–39). Although mice are often not considered to develop latent infection with *M. tuberculosis*, the model used in this study produces a stable, chronic paucibacillary lung infection that is ≤4 log_10_ CFU/lung (32–34, 40). Thus, this model can be used to evaluate the activity of drugs and regimens against an overall non-replicating *M. tuberculosis* population representing the presumed upper limit of infection in human LTBI. Moreover, this model has been validated through a number of experiments demonstrating the equivalent efficacy of the three rifamycin-based LTBI regimens currently recommended by the WHO (32–34, 41), as well as the 1-month daily isoniazid-rifapentine regimen recently evaluated in a phase 3 clinical trial (8). Furthermore, the superiority of these rifamycin-based regimens to six months of isoniazid in this model is consistent with the results of a meta-analysis of clinical trials suggesting that rifamycin-based regimens may have superior efficacy (42). Thus, experimental and clinical evidence supports the use of this model for the preclinical evaluation of LTBI treatment regimens.

In the present experiment, our *M. tuberculosis* challenge infection resulted in a larger infectious dose than expected. Our goal was to implant approximately 1.0-1.5 log_10_ CFU/lung (32–34), but our actual implantation was slightly more than 2 log_10_ CFU/lung. As a result, the *M. tuberculosis* lung burden was somewhat higher than intended at the start of treatment (Day 0), averaging 4.75 log_10_ CFU (Table 3). Nevertheless, the *M. tuberculosis* lung burden remained stable in untreated control mice throughout the duration of the experiment, indicating that the model was still suitable for the evaluation of LTBI regimens. Indeed, the bactericidal activity of the R_10_ (5/7), H_50_P_15_ (1/7), and B_25_ (5/7) control regimens, which resulted in decreases of 3, 4.5, and 4.9 log_10_ CFU/lung, respectively, after 12 weeks of treatment, was of the same magnitude as that observed in previous studies (32–34). Thus, the higher implantation and Day 0 CFU counts did not affect the relative activity of the drugs against this stable bacterial population in the mouse lungs.

One of the most striking findings from this study was the apparent duration of bactericidal activity associated with a single dose of the long-acting bedaquiline formulation. One injection of B_LAI-160_ at Day 0 continued to exert bactericidal activity up to the 12-week time point; these data are supported by the PK data, indicating that the plasma bedaquiline levels remained above the MIC for *M. tuberculosis* for at least 12 weeks post-administration. Additional studies with longer follow-up after dosing are clearly needed to understand the full extent of activity of the LAI formulation. When considering this enduring bactericidal activity, it is possible that just two injections of the long-acting bedaquiline formulation, spaced four weeks apart, could be as active as any of the WHO-recommended LTBI treatment regimens. This is a timely finding, as the recently reported success of the 1-month daily isoniazid-rifapentine regimen (8) may have set a new bar for a short-course LTBI regimen. Even more promising is the idea of combining a single injection of the long-acting bedaquiline formulation with a compatible one- or two-week oral regimen. Such a further decrease in patient-provider encounters could revolutionize LTBI treatment. In addition, our results suggest that an LAI formulation of bedaquiline could meet most, if not all, criteria in a recently proposed target product profile for LTBI treatment using LAI formulations, including many of the criteria for an “ideal” regimen (9). Finally, currently recommended short-course regimens for LTBI, as well as the newly reported 1-month isoniazid-rifapentine regimen, all contain isoniazid and/or a rifamycin (2, 8) and are therefore not expected to be effective against LTBI caused by MDR *M. tuberculosis*. The present results further support bedaquiline-based regimens for LTBI treatment in contacts of patients with MDR-TB (43), and LAI formulations in particular could significantly simplify what could be an otherwise long and complicated treatment.

## METHODS

### Long-acting bedaquiline formulation

A long-acting formulation of bedaquiline was developed as a microsuspension containing bedaquiline and D-tocopherol polyethylene glycol 1000 succinate in a ratio of 4:1 and 50 mg/mL of mannitol. The concentration of bedaquiline in the final formulation was 200 mg/mL.

### PK studies

The mouse PK procedures were approved by the local Johnson & Johnson Ethical Committee. Male Swiss mice (4-5 weeks old) were purchased from the Janvier Breeding Center (Le Genest Saint-Isle, France). All animals were housed under controlled conditions (specific pathogen free, 23°C, 60% humidity, and normal light-dark cycle) and had access to food and water *ad libitum*. A single, 160 mg/kg dose of long-acting bedaquiline was administered by intramuscular injection to five mice. Blood samples were taken from each animal at 1, 4, 7, 24, 168, 336, 504, 672, 840, 1176, 1512, 1848 and 2184 hours after injection. Within 1 hour of sampling, the blood samples were centrifuged. After centrifugation, plasma was collected and stored at −18 °C. At all times, blood/plasma samples were protected from light and placed on melting ice. To measure bedaquiline and its M2 metabolite in plasma, all samples were analyzed using a qualified LC-MS/MS method (44). The samples were subjected to a selective sample cleanup, followed by LC-MS/MS. Samples were quantified against calibration curves prepared to cover the concentration range of the study samples. The curves were prepared in the same matrix as the study samples. For each analytical batch, independent quality control samples, prepared in the same matrix as the samples, were analyzed together with the study samples and calibration curve. Individual plasma concentration-time profiles were subjected to a non-compartmental analysis using the linear up/log down trapezoidal rule for all data. Peak plasma concentrations (C_max_), corresponding peak times (T_max_), and the area under the plasma concentration-time curve from time zero to time t (AUC_0-t_), where t is the sampling time corresponding to the last measurable concentration above the limit of quantification (5 ng/mL), and from time zero to infinity (AUC_0-∞_), were calculated.

### Mycobacterial strains

*M. bovis* rBCG30, a recombinant strain in the Tice BCG background that overexpresses the *M. tuberculosis* 30-kilodalton major secretory protein (45), originally provided by Professor Marcus A. Horwitz, and *M. tuberculosis* H37Rv, American Type Culture Collection strain 27294, were separately mouse-passaged and frozen in aliquots. Frozen stocks were thawed and grown in liquid culture media to an optical density at 600 nm of around 1.0, and the actively growing cultures were diluted for use in experiments as follows: 10-fold in assay media to prepare MIC assay inoculum (H37Rv only), and 50-fold (rBCG30) or 100-fold (H37Rv) in phosphate-buffered saline to prepare suspensions used for infections.

### Media

Liquid culture medium was Middlebrook 7H9 broth supplemented with 10% (v/v) oleic acid-albumin-dextrose-catalase (OADC) enrichment, 0.5% (v/v) glycerol, and 0.1% (v/v) Tween 80. Assay medium for MIC determination was 7H9 broth supplemented with 10% (v/v) OADC and 0.5% (v/v) glycerol but without Tween 80. All plating was done on 7H11 agar supplemented with 10% (v/v) OADC enrichment and 0.5% (v/v) glycerol. Lung homogenates (and their cognate tenfold dilutions) were plated on selective 7H11 agar (7H11 agar containing 50 μg/mL carbenicillin, 10 μg/mL polymyxin B, 20 μg/mL trimethoprim, and 50 μg/mL cycloheximide) (46) that was further supplemented with 0.4% activated charcoal to adsorb any drug carried over in the homogenates (28). For differentiating *M. bovis* rBCG30 from *M. tuberculosis*, selective 7H11 agar was additionally supplemented with either 40 μg/mL hygromycin B, selective for *M. bovis* rBCG30 and not *M. tuberculosis*, or 2-thiophenecarboxylic acid hydrazide (TCH), selective for *M. tuberculosis* and not *M. bovis*, at 4 and 200 μg/mL in non-charcoal-containing and charcoal-containing agar, respectively. Difco Middlebrook 7H9 broth powder, Difco Mycobacteria 7H11 agar powder, and BBL Middlebrook OADC enrichment were obtained from Becton, Dickinson and Company. Glycerol and Tween 80 were obtained from Fisher Scientific, and activated charcoal was obtained from J.T. Baker. All selective drugs were obtained from Sigma-Aldrich/Millipore Sigma.

### MIC assays

MICs of rifampin, isoniazid, rifapentine, and bedaquiline for our *M. tuberculosis* H37Rv stock strain were previously determined using the broth macrodilution method and are 0.25, 0.03, 0. 03-0.06, and 0.06 μg/mL, respectively (28, 35, 47). The same broth macrodilution method was used to compare the MIC of the long-acting bedaquiline formulation with orally-administered bedaquiline; all bedaquiline was provided by Janssen. *M. tuberculosis* H37Rv was inoculated into polystyrene tubes containing 2.5 mL assay broth (at 5 log_10_ CFU/mL) containing bedaquiline concentrations ranging (by two-fold serial dilutions) from 64 to 0.0039 μg/mL. The MIC was defined as the lowest concentration that inhibited visible bacterial growth after 14 days of incubation at 37 °C.

### LTBI mouse model

The mouse model procedures were approved by the Johns Hopkins University Animal Care and Use Committee. The paucibacillary mouse model used in this study was previously described (33, 34). Female BALB/c mice (n = 150), aged 10 weeks were purchased from Charles River Laboratories. Mice were housed in individually ventilated cages (up to five mice per cage) with sterile wood shavings for bedding and with access to food and water *ad libitum*. Room temperature was maintained at 22-24 °C with a 12 h light/dark cycle. At each time point, mice were sacrificed by intentional isoflurane overdose by inhalation (drop method) followed by cervical dislocation. Mice were immunized by aerosol infection with a nebulized suspension of *M. bovis* rBCG30 using a Glas-Col Full-Size Inhalation Exposure System, per the manufacturer’s instruction; the day after infection, five mice were sacrificed to determine *M. bovis* rBCG30 lung implantation. Six weeks after the immunizing infection, mice were challenged by aerosol infection with a nebulized suspension of *M. tuberculosis* H37Rv, and the day after infection, five mice were sacrificed to determine both the *M. tuberculosis* lung implantation and the level of *M. bovis* rBCG30 lung infection. The bacterial concentrations of the suspensions used for infections and the subsequent lung implantation CFU counts were determined as previously described (34, 48) and as outlined in **Tables S1** and **S2**, respectively.

### Treatment

Treatment was initiated on Day 0, thirteen weeks after *M. tuberculosis* challenge infection. This was about twice as long as the usual incubation period but was necessitated by a delay in the availability of the LAI formulation. Five mice were sacrificed on Day 0 to determine pretreatment bacterial lung levels as previously described (34) and as outlined in **Table S3**. Mice were randomized into one of the ten treatment regimens described in Table 2. Doses of rifampin, isoniazid, and rifapentine were chosen to achieve similar plasma exposures (based on area under the plasma concentration-time curve) in mice as are achieved with recommended human doses for treatment of LTBI (49, 50). The daily bedaquiline dose of 25 mg/kg represents the standard dose used in TB treatment studies in mice (22, 28). The daily oral bedaquiline doses of 2.67, 5.33, and 8 mg/kg were chosen to administer the same total amount of bedaquiline over a 12-week period as one, two, and three doses, respectively, of the LAI bedaquiline formulation, which was administered at 160 mg/kg per dose. For drugs administered by gavage (all drugs except for the long-acting bedaquiline formulation), drug solutions were prepared to deliver the desired dose based on an average mouse body mass of 20 g in a volume of 0.2 mL; drug solutions were prepared weekly and stored at 4 °C. Rifampin and isoniazid were purchased from Millipore-Sigma and prepared in distilled water; rifapentine (Priftin®) tablets were purchased from a local pharmacy and prepared in distilled water. Orally-administered bedaquiline was dissolved in 20% (w/v) 2-hydroxypropyl-β-cyclodextrin adjusted to pH ~2 with 1N HCl. The LAI bedaquiline formulation was stored at 4 °C. The 160 mg/kg dose (based on an average mouse body mass of 20 g) was administered by two intramuscular injections (8 μL each, one into each hind thigh) using a BD Veo™ insulin syringe with BD Ultra-Fine™ 3/10 mL 6 mm × 31 G needle with half-unit scale. Treatment was administered for up to 12 weeks, with five mice per treatment group sacrificed 4, 8, and 12 weeks after Day 0. At sacrifice, lungs were removed and homogenized, and CFU counts were determined as previously described (34) and as outlined in **Tables S4-S7**. At each time point, the bacterial burden of *M. bovis* rBCG30 was calculated using the CFU/lung values determined on hygromycin-containing agar. The *M. tuberculosis* implantation was calculated using the CFU/lung values determined on TCH-containing agar. The *M. tuberculosis* bacterial burden at Day 0 and Week 4, 8, and 12 were calculated by subtracting the CFU/lung determined on hygromycin-containing agar from the total CFU/lung determined on plain agar. The primary outcome was the difference (decline) in *M. tuberculosis* lung CFU counts during treatment, compared to counts in the lungs of rBCG30-immunized but untreated negative control mice.

### Statistical analyses

CFU counts (*x*) were log-transformed as (*x* + 1) before analysis. The bactericidal activity of different treatment regimens at each time point was compared using one-way analysis of variance with Bonferroni’s correction for multiple comparisons. Analyses were performed using GraphPad Prism version 7.02.

## ACKNOWLEDGEMENTS

This work was funded by Janssen.

